# Bacterial community shift in nutrient-treated oil-bearing sandstones from the subsurface strata of an onshore oil reservoir and its potential use in Microbial Enhanced Oil Recovery

**DOI:** 10.1101/322891

**Authors:** Thanachai Phetcharat, Pinan Dawkrajai, Thararat Chitov, Pisanu Wongpornchai, Schradh Saenton, Wuttichai Mhuantong, Pattanop Kanokratana, Verawat Champreda, Sakunnee Bovonsombut

## Abstract

Microbial Enhanced Oil Recovery (MEOR) is a promising strategy to improve recovery of residual oil in reservoirs, which can be performed by promoting specific indigenous microorganisms. In this study, bacterial communities and the effects of elemental nutrient treatment of oil-bearing sandstone cores originated from six oil wells of an onshore reservoir was determined by tagged 16S rRNA gene amplicon sequencing, using Ion Torrent Metagenomic Sequencing Analysis. A total number of sequences were taxonomically classified into 43 phyla, 320 families, and 584 genera, with the dominant bacterial populations being related to *Deinococcus-Thermus*, and *Betaproteobacteria*. The nutrient treatment resulted in markedly increase in the relative abundance of *Gammaproteobacteria. Thermus, Acinetobacter*, and *Pseudomonas* were the most abundant genera. To our knowledge, this is the first report on the effect of elemental nutrients on alteration of bacteria communities attached to the oil-bearing rock. It provides comprehensive data on bacterial, physical, and chemical structures within a reservoir and demonstrates how these parameters can be co-analyzed to serve as a basis for designing a MEOR process. It also provides a model of how a bacterial community in reservoirs’ strata can be altered by nutrient treatment to enhance the efficiency of MEOR applications.

## 1. Introduction

Petroleum resources are becoming more limited, and while other alternative energy resources are being explored, Enhanced Oil Recovery (EOR) technologies are increasingly important. These technologies have been designed in response to the facst that the conventional petroleum recovery procedures can only retrieve 10 to 45 percent of crude oil (Denbina *et al.* 1991). Conventional EOR processes require the use of chemical or thermal methods or miscible gas injection. Some of the factors involved in these processes are harmful to the environment (Sen 2008). An EOR method that employs microorganisms, known as Microbial Enhanced Oil Recovery (MEOR), offers an alternative way to retrieve residual oil, especially from reservoirs with decreasing productivity, while being environmentally friendly (Sen 2008). MEOR has been estimated to increase recovery by up to one-third of the oil initially recovered in an oil reservoir (Kaster *et al.* 2012). This technology also requires comparatively low amounts of energy to operate and is economical (Lazar *et al.* 2007).

Many subsurface microorganisms have been reported to play crucial roles in MEOR. These include *Bacillus*, *Methanobacterium*, and *Clostridium* (Donaldson *et al.* 1989). Some aerobic bacteria, especially *Pseudomonas*, were also found in subsurface strata and often in association with the presence of nitrate. These microbes can function in MEOR through different mechanisms, including the production of gases (CO_2_, CH_4_, H_2_, and N_2_), low molecular weight acids, solvents (especially ethanol and acetone), biosurfactants, and biopolymers. These microbial metabolites can lead to increases in the efficiency of oil recovery by viscosity reduction, permeability alteration, and emulsification of crude oil. The microbial metabolites can decrease the surface and interfacial tensions and alter wettability, which results in increasing pore scale displacement (Kaster *et al.* 2012). One approach for MEOR application is to supply effective microorganism(s) or microbial metabolites from external sources to oil reservoirs (Sen 2008). Alternatively, MEOR can employ indigenous microbes that are supplied with additional nutrients to increase their metabolic activities which will enhance oil recovery. For this second approach, microbial communities in the subsurface strata, both the original and the new ones formed after alteration of subsurface conditions, are crucial keys to MEOR success.

Several studies have reported diversity of microbial communities in onshore and offshore oil reservoirs as well as the *in situ* and *in vitro* effects of MEOR on microbial community structures, using culture-dependent methods (Agrawal *et al.* 2010; Ghojavand *et al.* 2008). However, with the culture-dependent methods, a large portion of unculturable microbes that is indigenous to specific environments such as petroleum reservoirs (Donaldson *et al.* 1989). The culture-independent methods provide more comprehensive data for MEOR microbial communities, such as those carried out at Shengli oil field, China (Wang *et al.* 2014), and the offshore reservoirs in the Norwegian Sea, Norway (Kotlar *et al.* 2011). Most investigations of microbial communities within reservoir, however, have been performed using reservoir fluids. There have been very limited studies on microbial communities in oil reservoirs’ rocks. Because most microbes residing in oil reservoirs are attached to the subsurface strata (Brown *et al.* 2000), microbial community data obtained from the stratal rock samples would be more relevant to MEOR than those obtained from the reservoir fluids or production fluids. Indigenous microbial communities, moreover, could respond to extrinsic nutrient addition in such a way to result in enhanced oil recovery (Brown *et al.* 2000; Brown 2010). Inconsistencies have been observed in previous studies (Brown 2010) and often these studies lack comprehensive data on the dynamics of microbial communities resulting from the nutrient addition.

With these gaps in MEOR studies, we therefore aimed to conduct an investigation on the diversity of bacteria derived from the core rock regions of a reservoir using Ion Torrent Metagenomic Sequencing Analysis, to demonstrate how an *in vitro* nutrient treatment could affect bacterial community structures or cause a shift in bacterial communities, and to analyze the correlation between some key characteristics of the reservoir and its indigenous bacterial communities. These investigations were not only intended to provide a basis for the future *in situ* MEOR implementation at the study site (Mae Soon Reservoir, Fang Basin, Thailand), but also to demonstrate how the bacterial structure and the physical and chemical characteristics of a reservoir are related and how these parameters can be co-analyzed and manipulated, in order to serve as a basis for designing a MEOR process.

## 2. Materials and Methods

### 2.1 Core sample collection

Core samples used in this study were obtained from the reference core collection of Mae Soon reservoir of Fang oil field, Chiang Mai, Thailand (19°50′N 99°09’E) (Fig 1). This reservoir has been in production for more than 50 years, with average productivity of 900 barrels of oil per day (bbl/d), and is currently experiencing a decline in its production rate. Six oil-sand samples were aseptically drawn from the six-foot-long cylindrical cores from 6 oil wells in the reservoir. These oil-sand samples were from depths of 1,999-2,500 feet below the earth’s surface. The samples were aseptically hammered to obtain small pieces or course powder for further analysis.

**Figure 1.**
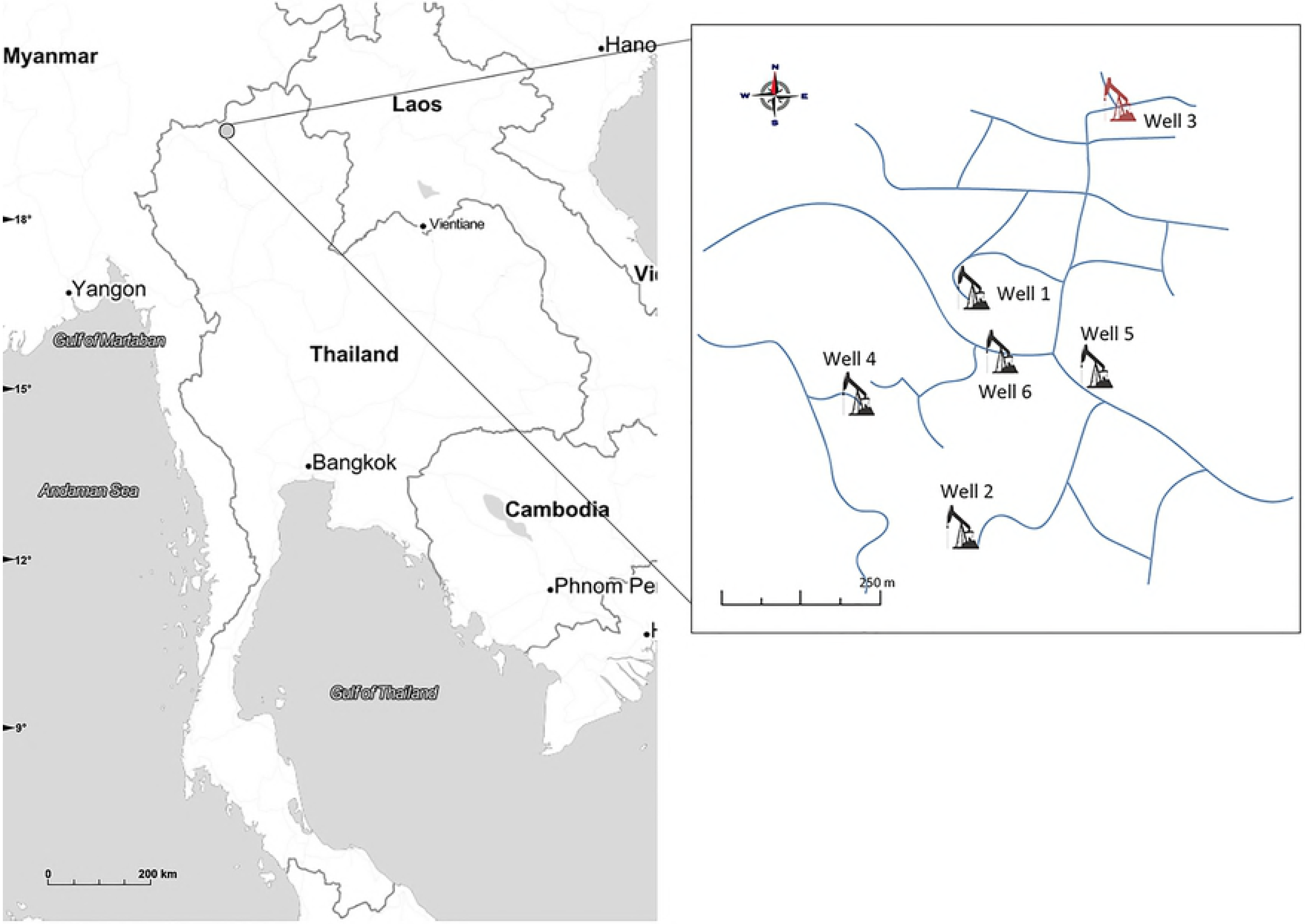
Location and distribution of six oil wells in Mae Soon reservoir, Fang oil field.

### 2.2 Reservoir characterisation

Temperatures and depths of oil reservoirs were retrieved from the well logs. Porosity, permeability and grain density of drilled core samples were determined as part of the routine core analysis (performed by GENLabs, Thailand). The lithology of core rock was analysed using an X-Ray fluorescence (XRF) spectrometer.

Produced water from the production wells was analysed for pH and salinity using a standard pH meter (Ohaus starter3100, USA) and an electrochemical analyser (Consort C933, Belgium), respectively. Concentrations of sulfate and nitrate in the produced water were determined using ion chromatography (Dionex ICS-3000, USA). Iron content was analysed using inductively coupled plasma optical emission spectrometry (ICP-OES).

### 2.3 Elemental nutrient enrichment

To study the effect of elemental nutrients on the changes in microbial composition within the oil sand samples, each of the six oil sand samples was divided into two 10-gram portions. The first portion was placed in a 250-ml screwed-cap bottle, which was filled completely with produced water (60°C) from the oil well. The cap was then tightened to create an anoxic condition. This treatment was performed to resemble the condition in the formation rock layer where the core originated. The samples treated with the produced water (assigned C1P – C6P) were used as control samples. Another portion of each oil sand sample was flooded with the produced water supplemented with elemental nutrients (0.12% (w/v) KNO_3_ and 0.034% (w/v) NaH_2_PO_4_), based on a core flooding protocol previously reported to be effective for oil recovery (Brown 2010). The controls and the treatments were incubated at 60°C in a water bath (Julabo, Germany) for 32 weeks.

### 2.4 DNA isolation and tagged 16S rDNA gene sequencing

The enriched core samples were separated from the fluids by centrifugation at 4,766 g for 5 min. The genomic DNA was extracted from 10 grams of the samples using 10 ml of lysis buffer (100 mM Tris-HCl, 100 mM Na-EDTA, 100 mM NaPO_4_, pH 8.0, 1.5M NaCl and 1% (v/v) CTAB) and 100 μl of proteinase K (Zhou *et al.* 1996). The solution was shaken horizontally at 37°C for 40 min. After that, 1 ml of 20% (v/v) SDS was added, and the solution was further incubated at 65°C for 30 min. The mixture was then centrifuged at 4,766 g for 5 min at room temperature. The mixture was extracted with chloroform/isoamyl alcohol (24:1 v/v) and centrifuged at 4,766 g for 10 min. Genomic DNA was precipitated with 0.6 volume of isopropanol at room temperature for 15 min, and the pellet was collected after centrifugation at 13,088 g for 20 min. The DNA pellet was washed with cold 70% (v/v) ethanol and dried at room temperature.

The purified metagenomic DNA was used as a template for amplification of the partial 16S rRNA gene. Universal bacterial primers E785F (5′-GGATTAGATACCCTGGTAGTCC-3′) and E1081R (5′-CTCACGRCACGAGCTGACG-3′) attached with tagged barcode sequences (Meyer *et al.* 2008) were used for amplifying the V5 and V6 regions of prokaryotic 16S rRNA genes. Polymerase chain reactions were performed using Phusion High-Fidelity DNA polymerase (Thermo Scientific, Massachusetts, USA) on MyCycler thermocycler (Bio-Rad, Hercules, CA) under the following conditions: initial denaturation at 98°C for 30 sec, 25 cycles of denaturation at 98°C for 10 sec, annealing at 66°C for 30 sec, and extension at 72°C for 30 sec, with a final extension at 72°C for 5 min. The PCR products were analysed on a 2 % (w/v) agarose gel and purified using the GF-1 ambiclean kit (Vivantis, Malaysia). Then, all amplicon products were quantified using Nanodrop spectrophotometer (LabX, CA). The sequences were determined using the ION PGM™ platform (Life Technologies, CA, USA) following the manufacturer’s recommended protocols. All 16S rRNA gene sequences of the oil sand samples were deposited into the NCBI Sequence Read Archive (SRA) with the project accession number SRP071710.

### 2.5 Data analysis

The sequencing dataset was initially processed by removing low-quality score reads with a cutoff of 20 for Phred quality score. The sequences were demultiplexed into specific groups based on the barcode sequences and then trimmed off the tagged and primer sequences using QIIME (Quantitative Insights into Microbial Ecology) version 1.9.1 (Caporaso *et al.* 2010). The sequences were clustered into Operational Taxonomic Units (OTUs) for classified taxon and alpha diversity analysis. Chimeric sequences were detected and removed by UCHIME (Edgar *et al.* 2011), based on the referenced dataset from the Ribosomal Database Project (Cole *et al.* 2014). The remaining sequences were clustered through an open-reference method using UCLUST at 97% similarity (Edgar 2010). The representative sequence of each cluster was determined taxonomically based on the nomenclature from the Greengene database (McDonald *et al.* 2012). To compare the bacterial diversities among the samples, alpha diversity index including observed OTUs, Shannon-Weaver index, and the Chao1 richness estimator were calculated from the OTU table using a cutoff of 16,828 reads per sample. The differences in bacterial profiles were analysed using statistical analysis of taxonomic and functional profiles software (STAMP) (Parks et al. 2014). The correlation between microbial community and some physical and chemical factors of the reservoir was analysed using Spearman’s Correlation (Hauke and Kossowski 2011).

## 3. Results

### 3.1 Physical characteristics of the reservoir

The oil-bearing sand core samples of Mae Soon reservoir were retrieved from the rock layer with the depths ranging from 1999 to 2482 feet. A previous survey has shown that the formation pressure of the deepest oil-bearing zone of this reservoir was 1,082 psi (unpublished data, provided by Northern Petroleum Development Center). The physical characteristics of the sandstone cores, together with the physical properties of the oil wells from which the cores originated are summarized in Table 1. It can be seen that the lithological nature of the cores, their porosity and grain density were slightly different, but the degrees of permeability, which reflect the mobility of petroleum liquid, had a greater variation among the samples.

**Table 1.**
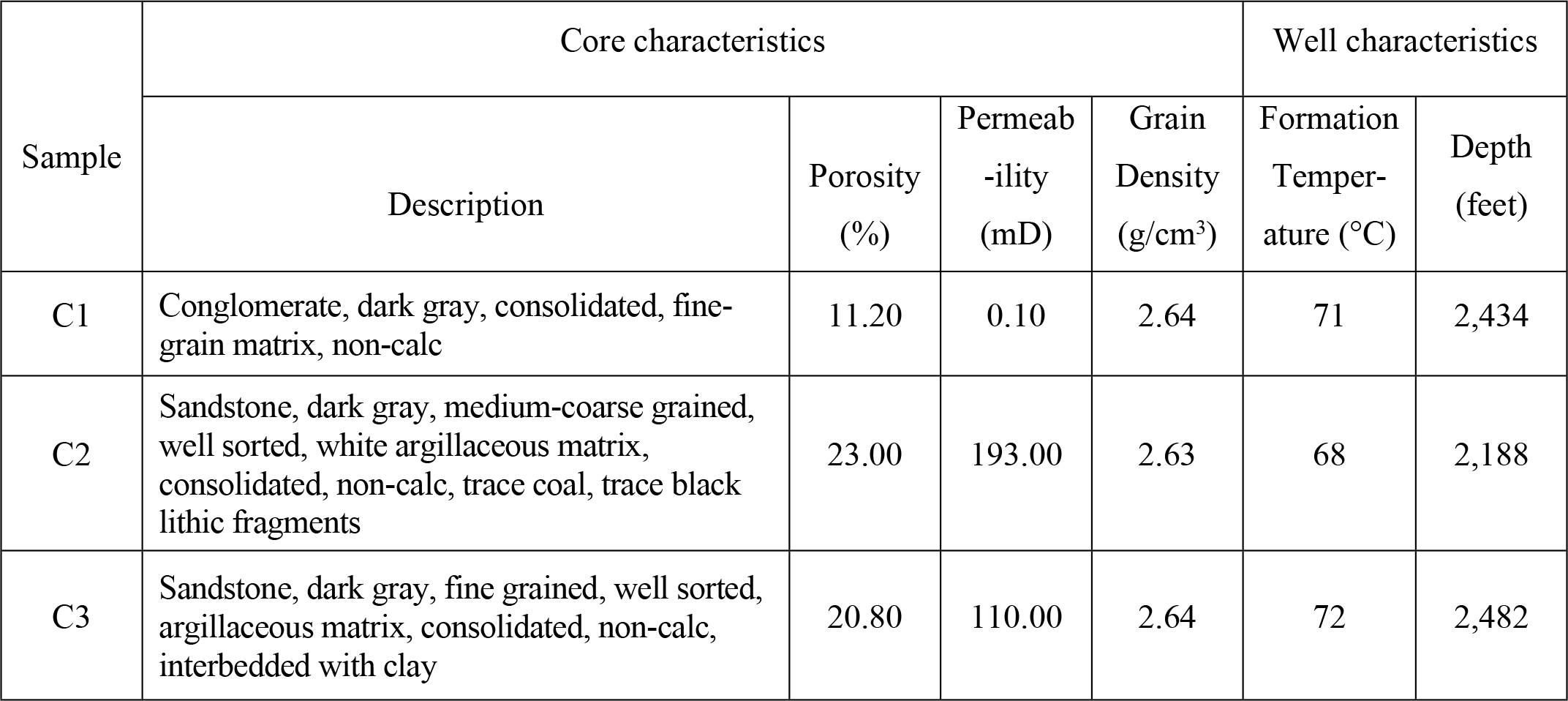
Characteristics of core samples and the wells from which core samples originated.

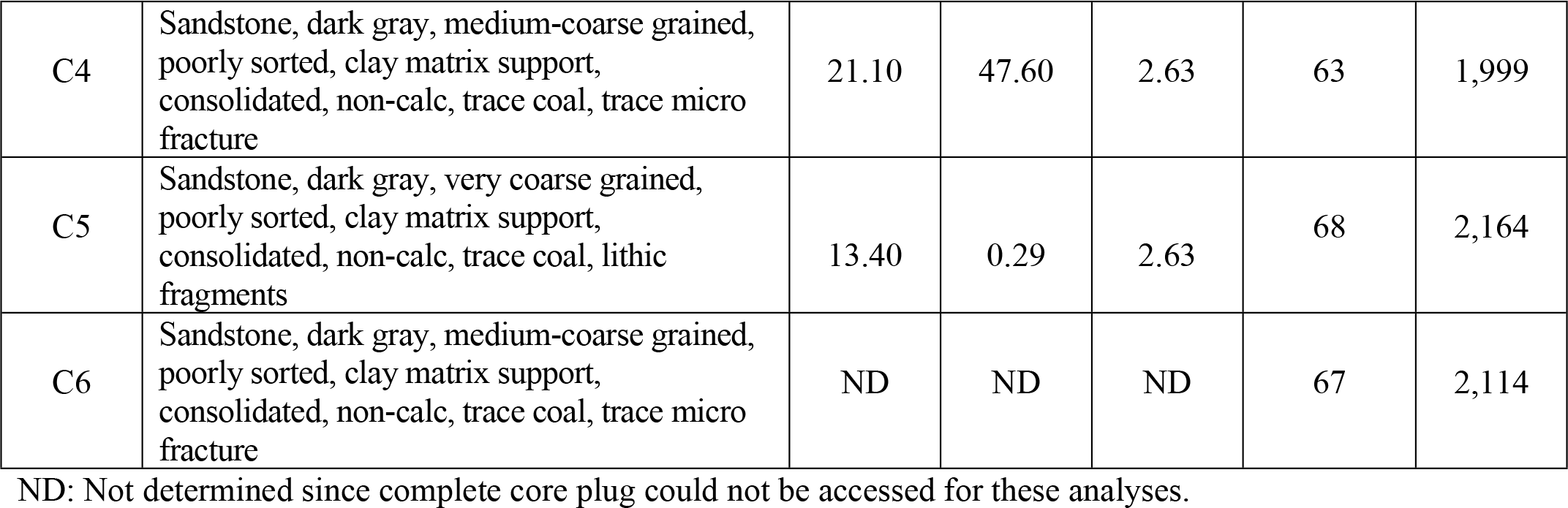

### 3.2 Chemical characteristics of the oil-bearing sandstone cores and produced water

The chemical properties of the oil-bearing sand cores and produced water were analysed in order to evaluate the possibility of applying a MEOR technique to the oil wells in this reservoir. Elemental compound analysis, using X-ray fluorescence spectrometer, revealed that the cores from this sandstone-based reservoir consisted mainly of SiO_2_ (80-85%). Elements including nickel, vanadium, rubidium, chromium, strontium, barium and zinc were present in all samples. The chemical properties of the cores are summarised in Tables 2 and 3.

**Table 2.** Percentages of elemental compounds of the core samples.

| Core sample | Compound proportion (%) |  |  |  |  |  |  |  |  |  |  |
| --- | --- | --- | --- | --- | --- | --- | --- | --- | --- | --- | --- |
|  | Al<sub>2</sub>O<sub>3</sub> | Fe<sub>2</sub>O<sub>3</sub> | K<sub>2</sub>O | MgO | MnO | Na<sub>2</sub>O | P<sub>2</sub>O<sub>5</sub> | SiO<sub>2</sub> | TiO<sub>2</sub> | CaO | Loss on ignition+SO<sub>3</sub> |
| C1 | 6.64 | 1.92 | 1.81 | 0.37 | 0.03 | 0.13 | 0.06 | 86.09 | 0.40 | 0.10 | 2.67 |
| C2 | 6.09 | 1.33 | 2.79 | 0.19 | 0.03 | 0.14 | 0.06 | 85.19 | 0.21 | 0.08 | 4.18 |
| C3 | 10.29 | 2.36 | 2.04 | 0.72 | 0.04 | 0.24 | 0.08 | 78.49 | 0.55 | 0.10 | 5.09 |
| C4 | 5.64 | 2.48 | 1.70 | 0.31 | 0.07 | 0.16 | 0.09 | 82.54 | 0.37 | 0.16 | 6.63 |
| C5 | 11.43 | 2.56 | 2.40 | 0.85 | 0.03 | 0.14 | 0.11 | 77.00 | 0.61 | 0.12 | 4.81 |
| C6 | 5.03 | 1.04 | 2.22 | 0.15 | 0.03 | 0.13 | 0.04 | 88.42 | 0.16 | 0.05 | 2.77 |

**Table 3.** The element contents of core samples.

| Core sample | Element content (ppm) |  |  |  |  |  |  |  |  |
| --- | --- | --- | --- | --- | --- | --- | --- | --- | --- |
|  | Ni | V | Rb | Y | Nb | Cr | Sr | Ba | Zr |
| C1 | 47.89 | 70.54 | 97.00 | 5.04 | ND | 136.35 | 31.07 | 155.43 | 194.60 |
| C2 | 36.10 | 44.87 | 147.75 | 2.99 | ND | 205.60 | 44.47 | 295.04 | 116.06 |
| C3 | 48.84 | 95.02 | 121.50 | 7.22 | 0.11 | 125.00 | 23.45 | 100.90 | 191.30 |
| C4 | 41.16 | 68.93 | 95.59 | 4.50 | ND | 165.14 | 23.32 | 103.55 | 214.38 |
| C5 | 53.23 | 110.18 | 133.03 | 7.96 | 1.89 | 143.38 | 41.08 | 141.40 | 238.69 |
| C6 | 30.16 | 32.20 | 117.61 | 4.58 | ND | 146.54 | 36.00 | 257.88 | 106.56 |

Note: ND - Not detected

Produced water from the oil wells was analysed for chemical properties, including pH and salinity. In addition, the concentrations of sulfate, nitrate, and iron, which can be the source of nutrition for microorganisms, were also determined. We found that the produced water from most wells (excluding well 3, which is presently abandoned) does not vary significantly in salinity, which is in the normal range of the salinity of onshore reservoirs. The pH of the produced water from the 6 wells tested showed that the wells were slightly alkaline. As for the elemental compounds, sulfate was not detected in the produced water from most wells, while nitrate was present in every well, and iron in 4 out of 5 wells (Table 4). An analysis of a representative crude oil sample from Mae Soon oil field showed that it had API gravity of 30°, the viscosity of 22 Cp (at 50 °C), and 18 (wt%) wax (unpublished data, provided by Northern Petroleum Development Center).

**Table 4.** The salinity and pH analysis and the dissolved compounds of produced water.

| Sample | Salinity (ppm) | pH | SO<sub>4</sub> (mg/L) | NO<sub>3</sub> (mg/L) | Fe (mg/L) |
| --- | --- | --- | --- | --- | --- |
| PW1 | 600 | 7.8 | ND | 0.76 | 0.10 |
| PW2 | 300 | 7.2 | 4.99 | 0.31 | 1.85 |
| PW4 | 500 | 8.6 | ND | 0.40 | ND |
| PW5 | 600 | 8.0 | ND | 0.34 | 0.46 |
| PW6 | 300 | 7.5 | ND | 0.24 | 1.94 |

Note: ND - Not detected.

### 3.3 Characterization of microbial diversity of the core samples

The purified DNAs from the sandstone core samples treated with produced water (C1P-C6P; control samples) and from those treated with produced water supplemented with elemental nutrient (C1N-C6N) were sequenced metagenomically. A total of 1,426,176 filtered reads were obtained from the ION PGM™ sequencing. The number of reads obtained from each sample ranged from 17,039 to 220,923 reads. The Operational Taxonomic Unit (OUT) in the samples was assigned from the filtered sequences at 95% similarity levels. The bacterial phylotypes ranged from 611 to 2,334 OTUs at the genetic distance of 0.05 with an average of 1,644 OTUs per sample. Rarefaction analysis was used to determine the microbial diversity in the filtered data set. The curves for nutrient-treated samples indicated that the population was less diverse than the control (Fig 2A). The highest bacterial richness was observed in sample C5N, which was in accordance with the diversity estimator, Chao 1. According to Shannon’s diversity indices, greater bacterial diversities were observed in the controls than in the nutrient-treated samples, with the exception of sample C4, in which bacterial diversity increased after being treated with elemental nutrient (Fig 2B). The diversity indices are summarised in Table 5.

**Figure 2.**
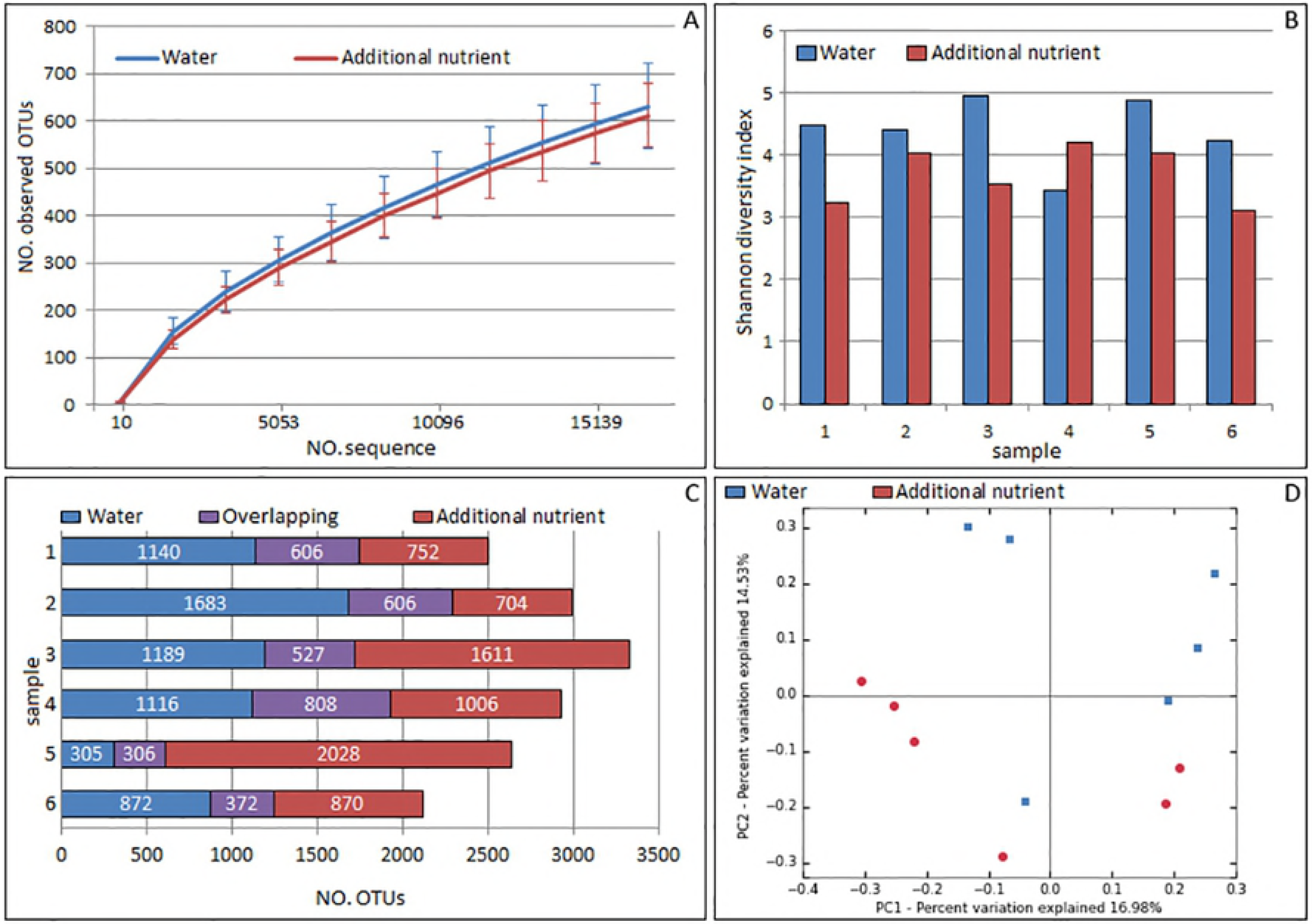
The dataset and diversity indices of metagenomic sequences. (A) Rarefaction curves of observed OTUs for the control and nutrient-treated groups, (B) Shannon diversity index comparison between the two treatments, (C) the number of unique and shared OTUs between the two treatments, (D) The principle coordinates analysis (PCoA) plot showing differences in bacterial community between the control and the nutrient-treated group.

**Table 5.**
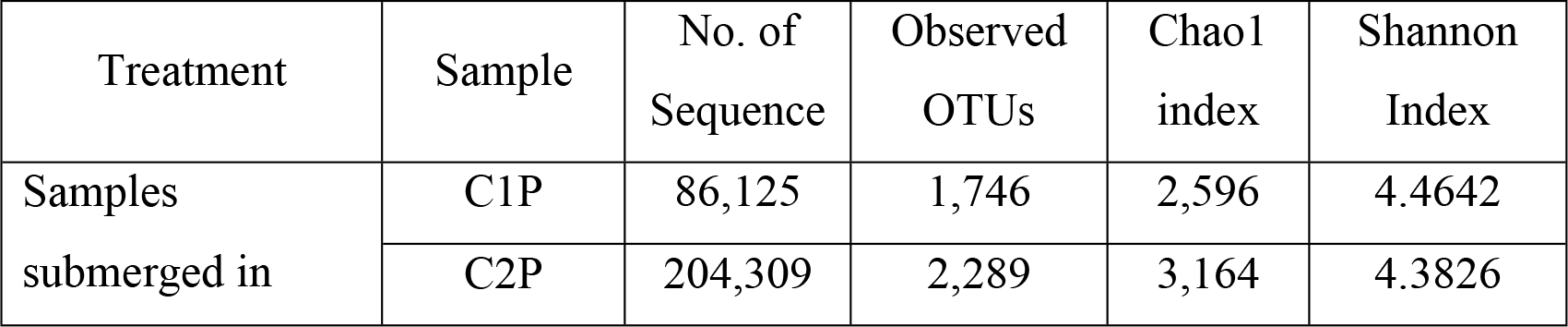
Number of sequences, OTUs, and alpha diversity indices of the bacterial community in the sand core samples treated with oil well produced water and produced water supplemented with elemental nutrients.

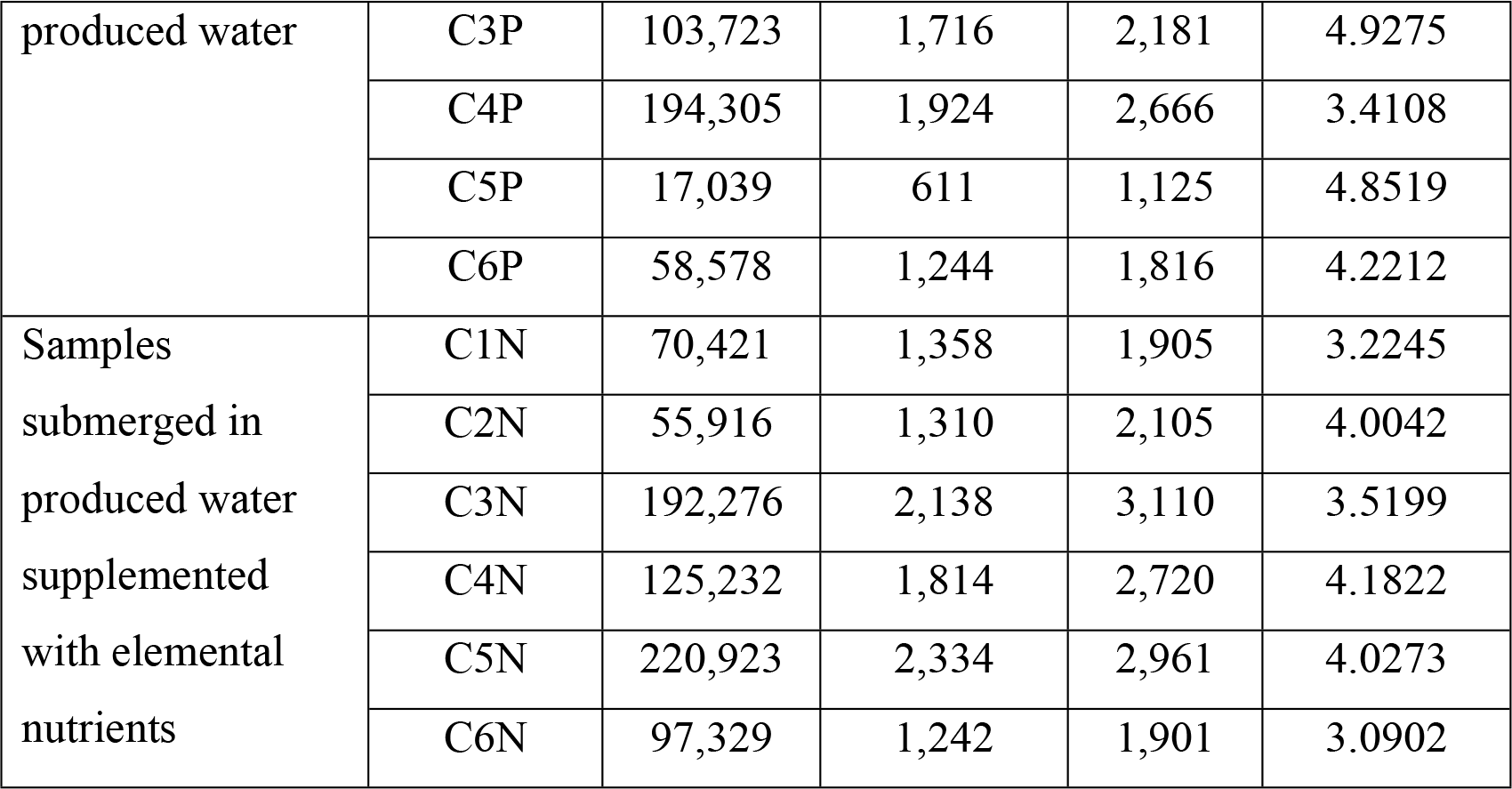

Bacterial communities were analysed in the oil-bearing part of the core samples submerged in the produced water and additional elemental nutrients. According to the shared OTUs chart (Fig 2C), bacterial groups in the sandstone core were altered after being treated with elemental nutrients. Smaller numbers of shared OTUs between the two treatments were observed compared to unique OTUs found in either produced water-treated or nutrient-treated samples. More than one thousand unique OTUs were found in produced water-treated samples C1P – C4P and nutrient-treated samples C3N – C5N. Principle Coordinated Analysis (PCoA) was performed to review the relationship of bacterial communities in the cores subjected to two treatments. PCoA indicated that the bacterial community profiles were associated with the treatment conditions used (Fig 2D). Comparison of the bacterial community under the different treatments using unweighted UniFrac (Lozupone and Knight 2005), beta diversity of bacteria as determined by permutational multivariate analysis of variance (PERMANOVA) was significantly different between the two treatments (P-value < 0.05).

### 3.4 Bacterial community structures in oil-bearing sandstone cores treated with produced water and additional elemental nutrients

Bacterial community profiles were studied between the two treatments at the phylum level (Fig 3). More diverse bacterial groups were observed among the control samples than in the nutrient-treated samples. In the controls, bacteria in phyla *Deinococcus-Thermus*, *Proteobacteria* (Beta), *Acidobacteria*, and *Firmicutes* (Clostridia) were detected in large proportions, which were 77.53% in C4P, 32.59% in C1P, 39.23% in C6P, and 31.81% in C5P. When the samples were treated with additional elemental nutrients, the bacterial groups became less diverse. The predominant phyla were mainly *Deinococcus-Thermus*, and *Proteobacteria* (Gamma). *Deinococcus-Thermus* was the major taxon found in C4N, C5N, and C2N (with proportions of 78.93%, 78.56%, and 66.30%, respectively), while *Proteobacteria* (Gamma) clearly dominated samples C1N, C6N, and C3N (with proportions of 81.67%, 77.30%, 60.26%, respectively).

**Figure 3.**
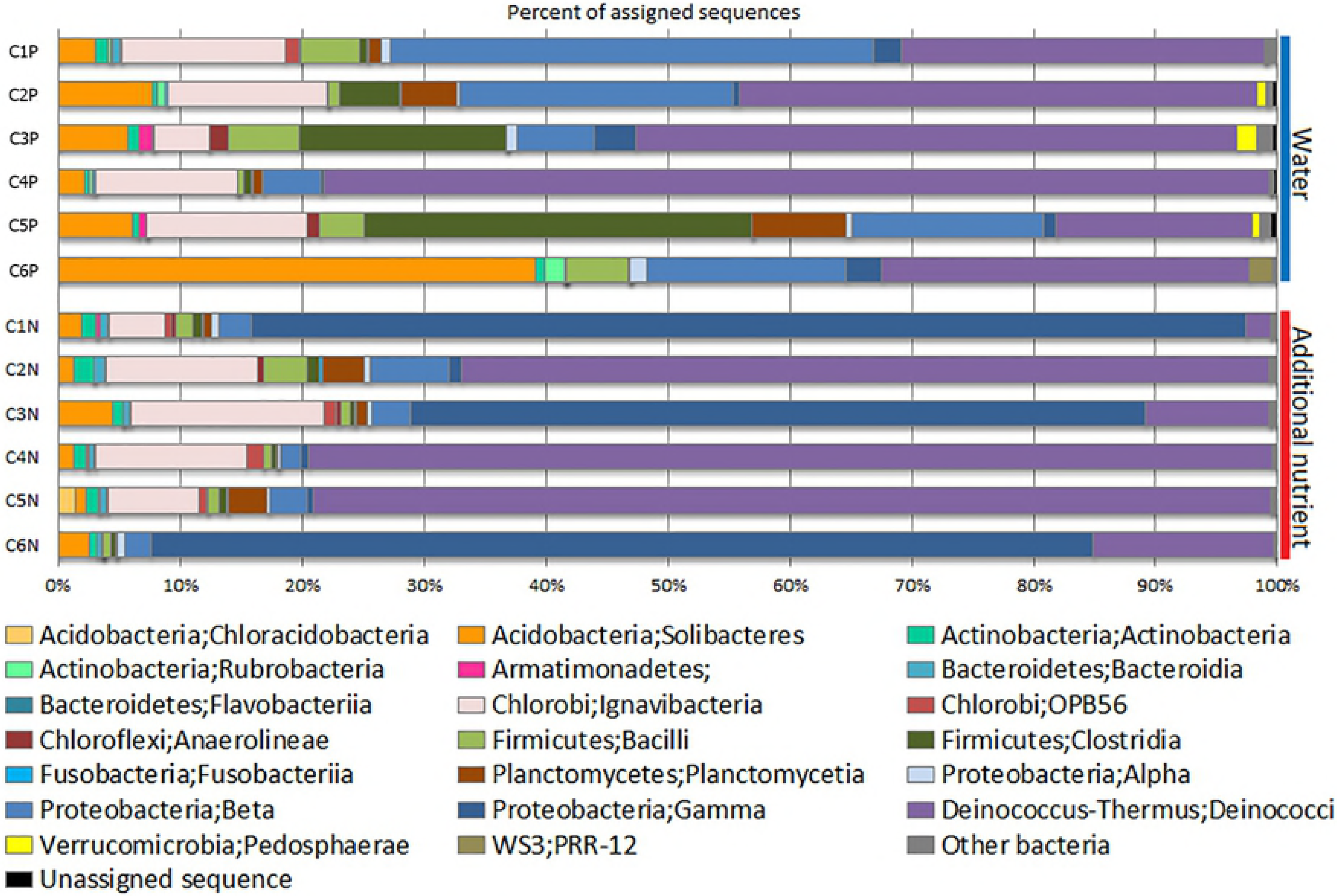
Taxonomic classification of OTUs at phylum level for the control and the nutrient-treated sandstone core samples.

The shifts in bacterial communities as a result of elemental nutrient treatments also resulted in decreases in proportions of some bacterial phyla. *Acidobacteria* and *Proteobacteria* (Beta) were the phyla that obviously decreased in every core sample after being treated with the elemental nutrient supplements. In addition, *Firmicutes* (Clostridia) also decreased in the majority of the nutrient-treated samples.

In the order level, the top five orders in the control group, according to the OTU abundances, were *Thermales* (in C4P, C3P, and C2P), *Solibacterales* (in C6P), *Burkholderiales* (in C1P and C2P), *Clostridiales* (in C5P), and SM1H02 (in C1P, C5P, and C2P). For the nutrient-treated group, the top five orders were *Pseudomonadales* (in C1N, C6N and C3N), *Thermales* (in C4N, C5N, and C2N), SM1H02 (in C3N, C2N, and C4N), *Solibacterales* (in C3N), and *Pirellulales* (in C2N and C5N), although the latter two orders had the abundances smaller than 10%.

We also determined the effects of different treatment conditions on bacterial diversity at the genus level. The abundances of bacterial genera were different between the samples submerged in produced water and those treated with elemental nutrient (Fig 4). In the overall picture of the reservoir, *Thermus* was the most abundant genus, both in the samples submerged in produced water and in the samples treated with additional nutrients, having average distributions of 33.79% and 39.97%, respectively. Treatment with the elemental nutrient caused a slight increase of this genus (6.18%). The populations of this genus increased in samples C2N (19.97% to 65.88%), C4N (74.46 to 78.13%), and C5N (5.15% to 76.79%); but decreased in samples C1N (27.30% to 0.55%), C3N (47.31% to 9.86%), and C6N (28.54% to 8.61%). Other bacterial genera that were obviously altered as a result of nutrient treatment were *Pseudomonas* and *Acinetobacter*. *Pseudomonas* increased up to 19.68%, while *Acinetobacter* increased up to 15.51%. Alterations of these genera were not consistent in all samples, although they increased in the majority of the samples. For examples, the relative abundance of *Pseudomonas* increased in nutrient-treated samples C1N, C3N, C4N, and C6N. *Acinetobacter* increased in C1N, C2N, C3N, C4N, and C6N. The most prominent increase in *Acinetobacter* was observed in nutrient-treated sample C1N (from 0.6% in C1P to 63.86% in C1N; increased by 63.25%), and *Pseudomonas* in C3N (from 0.93% to 53.77%; which was up 52.84%).

**Figure 4.**
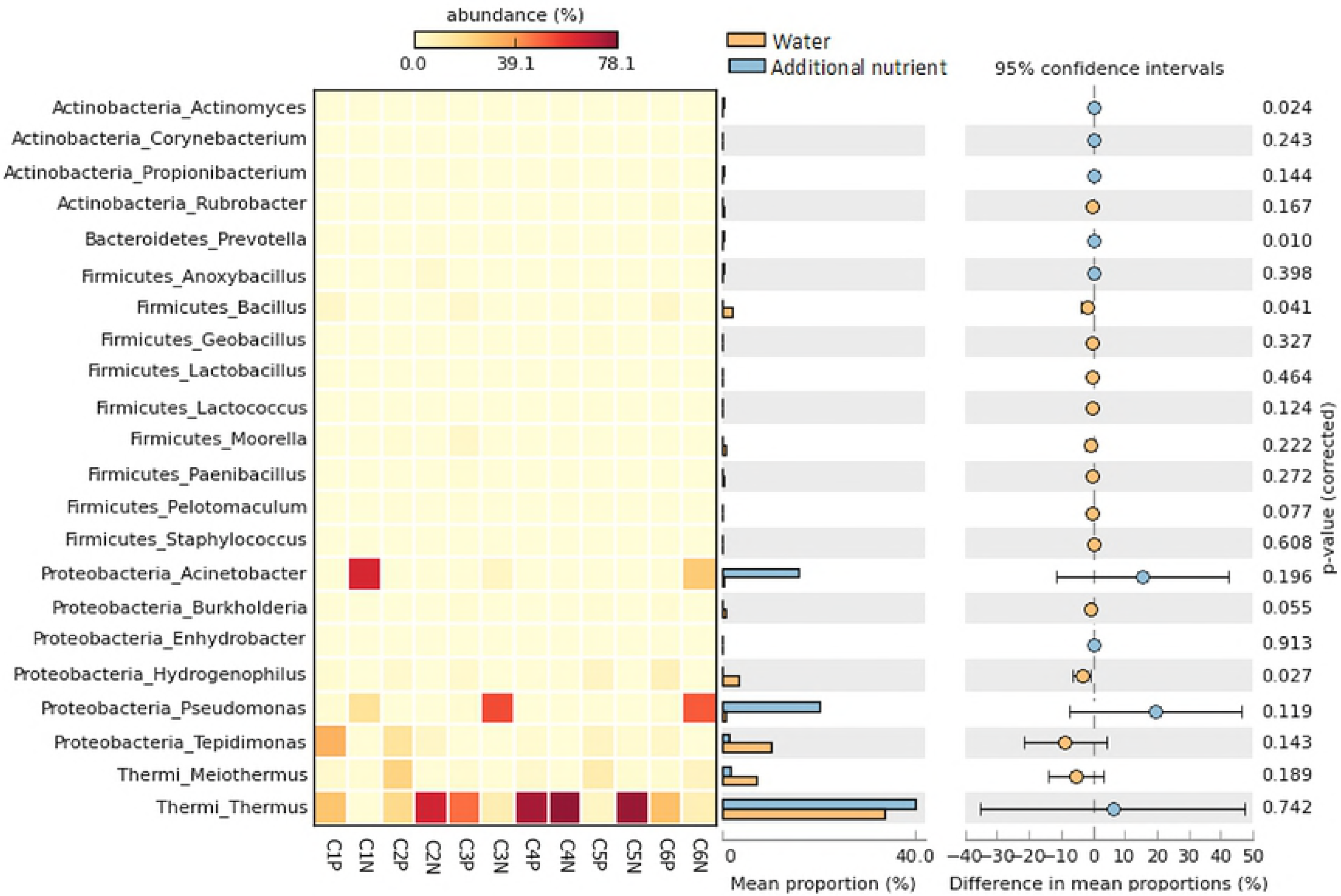
Comparative analysis of genus abundance in bacterial communities from the controls (C1P-C6P) and nutrient-treated samples (C1N-C6N).

On the other hand, some genera, such as *Tepidimonas, Meiotermus, Hydrogenophilus*, and *Bacillus*, experienced an average decrease in their proportions after nutrient treatment (Fig 4). The decrease of relative abundance of *Tepidimonas* decreased most obviously in the sample C1 (from 32.89% (C1P) to 0.55% (C1N)), *Meiothermus* in C2 (from 22.48% (C2P) to 0.42% (C2N)), *Hydrogenophilus* in C6 (from 7.94% (C6P) to 0.0010% (C6N)), and *Bacillus* in C6 (from 3.67% (C6P) to 0.08% (C6N)).

## 4. Discussions

### 4.1 Characteristics of oil reservoirs in relation to the potential for MEOR application

Microbial communities in different oil reservoirs and alteration in their microbial composition depend greatly on intrinsic conditions of the strata and the extrinsic factors influencing them. In this study, the oil-bearing sands from drilled cores of the Mae Soon reservoir in six wells in Fang oil field, situated in Chiang Mai in the north of Thailand, were characterised for their physical and chemical properties. These characteristics give us an understanding of the nature of this reservoir, its connection to its microbial communities, and the potential for application of Microbial Enhanced Oil Recovery (MEOR) technology (Lin *et al.* 2014).

Mae Soon reservoir is one oil-bearing reservoir in the Fang oil field, with the oil sand at a depth between 1,999 to 2,500 feet. The depth affects the temperature and pressure of the formation, which, in turn, can limit microbial growth and metabolism (Donaldson *et al.* 1989). Because Mae Soon reservoir is a shallow reservoir, the formation temperature is not too high (63 to 72°C) and this range of temperatures can support growth and metabolism of moderate thermophiles (Canganella and Wiegel 2014). The formation pressure of the deepest oil-bearing zone of Mae Soon is 1,082 psi, which should not affect microbial metabolism, according to Donaldson *et al.* (1989), who report that pressure lower than 1,400-2,800 psi does not limit bacterial growth.

Analysis of the sand core samples drawn from the reservoir showed that different sand core samples varied greatly in their degrees of permeability, which ranged from less than 1 to over 100 mD. Permeability of less than 75 mD is expected to limit effective microbial transport through the reservoir (Kalish *et al.* 1964). Thus, from the MEOR perspective, the permeabilities of core samples C2 and C3 seemed to be promising. As for the porosities, which reflect the fluids (which can be water or hydrocarbons) contained within the reservoir rock (Kalish *et al.* 1964), they were within the range found in most commercial petroleum reservoirs, ranging from 10-25%. This parameter might affect the cell transport in the pore space of the oil reservoir (Donaldson *et al.* 1989). Iglauer *et al.* (2010) observed that rocks with porosity about 20% supported oil recovery of additional incremental oil from sandstone core up to 75%, using the surfactant recovery method.

Analysis of the lithology of the rock, the cores from this sandstone-based reservoir, and this is favorable for biotechnological oil-recovery method (Brown *et al.* 2000). The cores contained elements commonly found in the other sandstone reservoirs’ strata (Olsson-Francis *et al.* 2016). The produced water, which was low in salinity and had neutral to slightly alkaline pH, could also support microbial growth (Donaldson *et al.* 1989). The crude oil (which, in Mae Soon reservoir, includes paraffin) could also be used by microorganisms as a carbon source. The oil gravity and its viscosity are also within the acceptable ranges for MEOR processes (Portwood 1995).

The physical and chemical properties of the oil-bearing rock and the properties of the produced water from the oil wells within a reservoir can determine the success of the MEOR method. The factors that are considered to support MEOR are: rock with a permeability greater than 75 mD, formation water with less than 10% NaCl and pH between 4-9, crude oil with API gravity greater than 18°, and formation temperatures of less than 75°C (Bryant and Burchfield 1989). According to these recommended conditions, this reservoir, in general terms, is favorable for MEOR application.

### 4.2 Bacterial community in oil-bearing sandstone cores

We have studied a microbial community in the Mae Soon oil-bearing sand reservoir. The oil-bearing sand from cores retrieved from six wells was placed under conditions resembling the reservoir’s conditions, flooded with the produced water in an anoxic condition at a raised temperature. We also investigated the microbial communities of the sand core samples from the reservoir after the addition of elemental nutrients. We did this to evaluate the impact of a change in extrinsic conditions on the bacterial communities, which will assist in designing a microbial enhanced oil recovery process.

The relative abundance of predominant taxa found in the oil-bearing sand core samples of Mae Soon reservoir showed that *Deinococcus-Thermus* (Deinococci) was the most abundant bacterial phylum in half (three out of six) of the samples tested. As for the other three samples, even though the *Deinococcus-Thermus* (Deinococci) was present, the predominant phyla differed. *Gammaproteobacteria* was predominant in one sample (C1P), while in samples, C5P and C6P, *Firmicutes* (Clostridia) and *Acidobacteria* (Solibacteres), were more abundant than others. This is interesting because *Gammaproteobacteria* had been found to be a more-common predominant bacterium in many other studies of oil-reservoir microbial communities than *Deinococcus-Thermus* (Wang *et al.* 2012). Thus, this reservoir’s bacterial communities being made up largely with *Deinococcus-Thermus* (Deinococci) is unique, although it is noted that the samples in other studies were mostly reservoir fluids. The microbial structure of the oil wells within the reservoir shared some common patterns, yet each well preserves its own microbial composition.

In the genus level, sequencing of 16S rRNA gene amplicons from the reservoir samples revealed several potential bacteria for microbial recovery process including *Thermus*, *Pseudomonas, Acinetobacter, Bacillus*, and *Clostridium* (Youssef *et al.* 2009). The most abundant genus was *Thermus*, which was found 27.30%, 19.97%, 47.31%, 74.46% and 28.54% in sample C1P, C2P, C3P, C4P, and C6P, respectively. This bacterium is generally found under elevated temperature conditions. It can grow at temperatures between 50 to 75°C (with the optimum temperature of 70°C), and is also found in high-temperature petroleum reservoirs (range 50 to 80°C) such as East Paris Basin (France) and Samotlor oil reservoir (Russia) (Bonch-Osmolovskaya *et al.* 2003; Jeanthon *et al.* 1995). Many thermophilic bacteria secrete biosurfactants; this mechanism supports the ability of hydrocarbon degradation by a microorganism (Shibulal *et al.* 2014).

Many facultative anaerobic bacteria found in the reservoir samples were bacteria that can potentially support the MEOR process. The *Bacillus* genus was found in sample C1P, C3P, and C6P. This genus occurred in the environment of many oil reservoirs (Biria *et al.* 2010). Some members of this genus are thermotolerant, capable of producing biosurfactants that play an important part in mobilising entrapped oil, and able to survive in extreme conditions. The characteristics and abilities of this genus are desirable in MEOR processes (Youssef *et al.* 2007). Besides these, *Clostridium* was also found, although not in a great abundance. This organism is one of the bacteria that can produce solvents and gasses under anaerobic conditions, which would support the efficiency of oil recovery by reducing viscosity (Sen 2008).

### 4.3 Bacterial community shift as a result of additional elemental nutrients

Microbial community in oil-bearing reservoirs is one of the key factors that contribute to the success of MEOR technology. In the effort of developing a MEOR process, researchers have studied microbial communities in petroleum fluids of oil reservoirs, such as formation water, injection water, and crude oil (Ren *et al.* 2011; Zhang *et al.* 2012). However, it has been noted that most bacteria occur in oil reservoirs attached to the strata and would not appear in formation fluids (Brown *et al.* 2000). Thus, the reservoir’s rock seems to be the better sample type than its petroleum fluid to reflect the bacterial community that may play a potential role in Enhanced Oil Recovery. Moreover, since bacterial composition in samples such as these can be influenced by factors in its surroundings (which, in this case, are mainly the oil trapped in the pores of the rock that can serve as a carbon source and the produced (formation) water which contains minerals or nutrients), bacterial composition in the oil-bearing strata can be altered, both in the natural situation and with intervention. We therefore investigated the bacterial community of the reservoir rock samples submerged in the reservoir’s produced water, which resemble their natural settings, and the shift in the bacterial community in the oil-bearing sandstone core samples treated with additional elemental nutrients, which can serve as a guideline for application of MEOR *in situ*.

Based on the sequencing results, the oil-bearing sandstone portion of the core samples harboured a great diversity of substrata microbes. The relative abundances of predominant taxa in the core samples altered after being treated with elemental nutrient.

*Deinococcus-Thermus* remained most abundant taxon in the reservoir’s community; comprised of 41.81%, which similar to the control group (40.88%). However, when considering the bacterial composition in each individual well, this phylum has significant changes. *Deinococcus-Thermus* increased in C2N, C4N, and C5N, with C5N being the sample in which this phylum increased most dramatically.

Obvious alteration of relative populations of bacteria as a result of nutrient treatment was not only observed with *Deinococcus-Thermus*, but also with *Gammaproteobacteria*. *Gammaproteobacteria* became predominant in three out of the six samples analysed, replacing the previously predominant *Betaproteobacteria, Deinococcus-Thermus*, and *Acidobacteria*, in C1P, C3P, and C6P, respectively. Interestingly, this bacterial taxon increased significantly, from 1.74% to 36.91% of the overall reservoir’s population. In contrary, *Proteobacteria* (Beta), *Acidobacteria* (Solibacteres), *Firmicutes* (Clostridia), and *Firmicutes* (Bacilli), were the taxa that experienced the decrease in overall population

Bacterial community profiles of nutrient-treated samples determined in this work were clearly different compared to the liquid phase of an offshore reservoir (Kotlar *et al.* 2011). According to Kotlar *et al.* (2011), Microbial communities in offshore reservoirs were dominated by *Delta/epsilon-Proteobacteria* (sulfate-reducing bacteria richness) and methanogens (mostly *Methanococcus* species). In our study, *Delta/epsilon-Proteobacteria* occurred in low abundance, while methanogens were not found. It is clear that bacterial distribution in the reservoir’s strata can greatly be influenced by the elemental nutrient supplement. The unique environment of an oil reservoir determines a specific frame of microbial communities. Therefore, it is necessary to investigate the microbial relative populations within a reservoir to provide information for supporting a MEOR process.

The bacterial populations that increased from elemental nutrient treatment observed in our work were mainly members of the genera *Thermus, Pseudomonas*, and *Acinetobacter*. A similar experiment has previously been done in production fluids, and the major microbes found in the enriched culture were bacteria in the order *Thermotogales*, methanogens related to *Methanobacterium* and *Methanococcus*, and sulfur-utilising archaea of the genus *Thermococcus* (Orphan *et al.* 2000). Nutrient treatment processes may give different results in bacterial community structures and patterns of distribution shifts, depending on the original seed culture, type of nutrient used, and conditions and time of incubation. Apart from these, the physical and chemical properties of the oil-bearing strata and the properties of produced water of reservoirs may have a significant contribution to the bacterial distribution patterns. To design a nutrient treatment process for a reservoir, which carries its own complexity, all these factors, therefore, must be taken into account.

### 4.4 Potential in MEOR application

MEOR processes employ microorganisms and/or their metabolic products that have the desired functions to enhance oil recovery (Youssef *et al.* 2009). *Deinococcus-Thermus* and *Proteobacteria* appeared as predominant phyla in the core rock samples from the Mae Soon oil reservoir in the Fang field.

Within the predominant phylum, which was *Deinococcus-Thermus*, *Thermus* was the most abundant genus, which is known as thermophilic hydrocarbon-degraders (Feitkenhauer *et al.* 2003) and was found to secrete rhamnolipids, a biosurfactant that can directly hydrolyse oil (Pantazaki *et al.* 2010). This genus converted crude oil into lighter hydrocarbons by decreasing aromatics, resins and asphaltenes in crude oil (Hao *et al.* 2004). It is, therefore, one of the promising thermophilic bacteria that can degrade hydrocarbons, which is one the desired characteristics for MEOR application.

*Betaproteobacteria* was found as the second most abundant group in the samples in the conditions resembling the *in situ* reservoir. Some of this bacterial group are acid-tolerant and can utilise various carbon substrates and can fix nitrogen gas (Dedysh 2011). So, in relation to MEOR, this group of bacteria, which was indigenous to the reservoir, may be useful even in its natural conditions. However, after the treatment of the oil-bearing sandstone core samples with the elemental nutrients used in this study, the *Proteobacteria* class that became predominant was *Gammaproteobacteria*, which became the second most abundant group in place of *Betaproteobacteria*. In terms of its benefit to Enhanced Oil Recovery, *Gammaproteobacteria* is considered one of the most promising groups that can be employed in MEOR processes. The most prominent genera belonging to *Gammaproteobacteria* class were *Pseudomonas* and *Acinetobacter*. These bacteria can produce many kinds of biosurfactant molecules that can lower interfacial tension of water, thus allowing mobilisation of bound hydrophobic molecules, which, in this case, is the oil that is trapped in the reservoir rock. The existence of *Pseudomonas* in onshore reservoirs had also been observed in several studies (Orphan *et al.* 2000; Wang *et al.* 2012).

Ren *et al.* (2011) showed that in their analysis of the 16S rRNA gene clone libraries, *Pseudomonas* was one of the predominant groups found in the produced water of Gudao oil reservoir, China. In another study, it was also found in the produced water samples originated from Daqing oil field, China (Hui *et al.* 2012), as the cultures recovered from a screening of thermophilic microorganisms. Fluid samples collected from producing wells of an offshore petroleum reservoir also contained a high ratio of *Pseudomonas* (Brakstad *et al.* 2008). The biosurfactants produced by this organism have a high potential for the recovery of crude oil (Nishanthi *et al.* 2010). Besides, *Pseudomonas* is known as a hydrocarbon-degrading bacterium, and it is effective in stabilising oil in water emulsion that can improve the ability of oil recovery (Banat *et al.* 2010). The fact that *Pseudomonas*, which is typically aerobic, was found in an oil reservoir and that it survived and became one of the organisms with greatest relative abundance after nitrate supplement was interesting. This genus could grow steadily in an anaerobic condition, as reported by (Kerschen *et al.* 2001), as it could use nitrate as a terminal electron acceptor for both assimilation and anaerobic respiration.

Another member of *Gammaproteobacteria* which was present in a large proportion was *Acinetobacter*. This is another potential bacterium for a MEOR process. Interestingly, previous studies have shown that this organism was predominant in the produced water samples of onshore Daqing oil field, and it was also found in the crude-oil-contaminated soil of Zhongyuan oil field, China (Hui *et al.* 2012; Zou *et al.* 2014). Several species of *Acinetobacter* produce substances involved in processes such as emulsification, interfacial tension reduction, and viscosity reduction (Al-Sulaimani *et al.* 2011), which are potentially useful for enhanced oil recovery. Metabolites from this bacterium have been used to increase displacement efficiency (Youssef *et al.* 2009).

Several bacterial genera such as *Clostridium*, *Klebsiella*, and *Bacillus*, known as potential producers of some metabolites that have the potential for enhancing oil recovery, were detected, although low in numbers. *Clostridium* and *Klebsiella* have been reported to increase the recovery process by reduction of oil viscosity (Sen 2008). *Bacillus* is among the organisms with high potential to produce biosurfactants and enzymes (Aurepatipan *et al.* 2017; El-Sheshtawy *et al.* 2015). Members of this genus could lower the interfacial tension of water, and thus could allow water to mobilise residual oil trapped in reservoirs’ oil-bearing zone. The spore-forming ability of *Bacillus* is also favourable for its survival in harsh conditions of different reservoirs (Youssef *et al.* 2007). This genus, however, was found in relatively low proportion in our study, even lower after the nutrient treatment. A treatment using elemental nutrients that could cause some bacterial genera, such as *Thermus, Pseudomonas*, and *Acinetobacter*, to prevail, obviously could not promote the growth of *Bacillus*. Alternatively, it could be possible that the organism remained in the dormant stage under the nutrient-treated conditions given in this study. Conditions that can elevate this genus to a higher relative abundance remain to be further investigated.

## 5. Conclusion

Bacterial community analysis of oil-bearing sandstone cores submerged in the produced water under high temperature conditions, revealed *Deinococcus-Thermus* (Deinococci) and *Betaproteobacteria* as the dominant groups. Bacterial community of the reservoir was altered after the elemental nutrients treatment, resulting in the shift of bacterial population, most obviously in *Betaproteobacteria* to *Gammaproteobacteria*. *Deinococcus-Thermus* remained the most abundant phylum. In the genus level, *Thermus, Acinetobacter*, and *Pseudomonas* were the most abundant. Members of these genera are capable of producing metabolites such as hydrocarbon-degrading enzymes and surfactants, which are potential for MEOR application. The microbial communities were supported by the reservoir’s physical and chemical characteristics, and thus may be promising for a MEOR process *in situ*. This work also serves as a guideline for evaluating the MEOR potential for other oil reservoirs.

## Acknowledgments

This work was financially supported by the National Research Council of Thailand (NRCT), the Environmental Science Research Center (ESRC), and the Graduate School, Chiang Mai University, Thailand. The authors would like to express our gratitude to the staff members of the Northern Petroleum Development Center, Defence Energy Department, for providing the samples used in this study and the accompanying data. We also would like to thank you, Dr. Robert Kieckhefer, guest lecturer, Petroleum Geophysics, Chiang Mai University, for his invaluable advice to improve our manuscript.

## References

1. Agrawal A, Vanbroekhoven K, Lal B. Diversity of culturable sulfidogenic bacteria in two oil-water separation tanks in the north-eastern oil fields of India. Anaerobe 2010;16:12–8.

2. Al-Sulaimani H, Joshi S, Al-Wahaibi Y et al. Microbial biotechnology for enhancing oil recovery: current developments and future prospects. Biotechnol Bioinformatics Bioeng 2011;1:147–58.

3. Aurepatipan N, Champreda, V, Kanokratana P et al. Assessment of bacterial communities and activities of thermotolerant enzymes produced by bacteria indigenous to oil-bearing sandstone cores for potential application in Enhanced Oil Recovery. J Pet Sci Eng 2017;163:295–302.

4. Banat IM, Franzetti A, Gandolfi I et al. Microbial biosurfactants production, applications and future potential. Appl Microbiol Biotechnol 2010;87:427–44.

5. Biria D, Maghsoudi E, Roostaazad R et al. Purification and characterization of a novel biosurfactant produced by Bacillus licheniformis MS3. World J Microbiol Biotechnol 2010;26:871–8.

6. Bonch-Osmolovskaya EA, Miroshnichenko ML, Lebedinsky AV et al. Radioisotopic, culture-based, and oligonucleotide microchip analyses of thermophilic microbial communities in a continental high-temperature petroleum reservoir. Appl Environ Microbiol 2003;69:6143–51.

7. Brakstad OG, Kotlar HK, Markussen S. Microbial communities of a complex high-temperature offshore petroleum reservoir. Int J Oil Gas Coal Tech 2008;1:211–28.

8. Brown L, Vadie A, Stephens J. Slowing production decline and extending the economic life of an oil field: new MEOR technology. SPE J 2000; DOI: 10.2523/59306-MS.

9. Brown L. Microbial enhanced oil recovery (MEOR). Curr Opin Microbiol 2010;13:316–20.

10. Bryant RS, Burchfield TE. Review of microbial technology for improving oil recovery. SPE Reservoir Eng 1989;4:151–4.

11. Canganella F, Wiegel J. Anaerobic thermophiles. Life 2014;4:77–104.

12. Caporaso JG, Kuczynski J, Stombaugh J et al. QIIME allows analysis of high-throughput community sequencing data. Nature Methods 2010;7:335–6.

13. Cole JR, Wang Q, Fish JA et al. Ribosomal Database Project: data and tools for high throughput rRNA analysis. Nucleic Acids Res 2014;42:633–42.

14. Dedysh SN. Cultivating uncultured bacteria from northern wetlands: knowledge gained and remaining gaps. Front Microbiol 2011;2:1–15.

15. Denbina E, Boberg T, Rottor M. Evaluation of key reservoir drive mechanisms in the early cycles of steam stimulation at Cold Lake. SPE Reservoir Eng 1991;6:207–11.

16. Donaldson EC, Chilingarian GV, Yen TF. Microbial enhanced oil recovery. Developments in Petroleum Science Vol. 22. Elsevier Science, 1989.

17. Edgar RC. Search and clustering orders of magnitude faster than BLAST. Bioinformatics 2010;26:2460–1.

18. Edgar RC, Haas BJ, Clemente JC et al. UCHIME improves sensitivity and speed of chimera detection. Bioinformatics 2011;27:2194–200.

19. El-Sheshtawy HS, Aiad I, Osman ME et al. Production of biosurfactant from Bacillus licheniformis for microbial enhanced oil recovery and inhibition the growth of sulfate reducing bacteria. Egyptian J Petrol 2015;24:155–62.

20. Feitkenhauer H, Müller R, MAuml H. Degradation of polycyclic aromatic hydrocarbons and long chain alkanes at 60-70 C by Thermus and Bacillus spp. Biodegradation 2003;14:367–72.

21. Ghojavand H, Vahabzadeh F, Mehranian M et al. Isolation of thermotolerant, halotolerant, facultative biosurfactant-producing bacteria. Appl Microbiol Biotechnol 2008;80:1073–85.

22. Hao R, Lu A, Wang G. Crude-oil-degrading thermophilic bacterium isolated from an oil field. Can J Microbiol 2004;50:175–82.

23. Hauke J, Kossowski T. Comparison of values of Pearson’s and Spearman’s correlation coefficients on the same sets of data. Quaestiones geographicae 2011;30:87–93.

24. Hui L, Ai M, Han S et al. Microbial diversity and functionally distinct groups in produced water from the Daqing Oilfield. China Petrol Sci 2012;9:469–84.

25. Iglauer S, Wu Y, Shuler P et al. New surfactant classes for enhanced oil recovery and their tertiary oil recovery potential. J Petrol Sci Eng 2010;71:23–9.

26. Jeanthon C, Reysenbach AL, L’Haridon S et al. Thermotoga subterranea sp. nov., a new thermophilic bacterium isolated from a continental oil reservoir. Arch Microbiol 1995;164:91–7.

27. Kalish P, Stewart J, Rogers W et al. The effect of bacteria on sandstone permeability. J Petrol Technol 1964;16:805–14.

28. Kaster KM, Hiorth A, Kjeilen-Eilertsen G et al. Mechanisms involved in microbially Enhanced Oil Recovery. Transport Porous Media 2012;91:59–79.

29. Kerschen E J, Irani VR, Hassett DJ et al. snr-1 gene is required for nitrate reduction in Pseudomonas aeruginosa PAO1. J Bacteriol 2001;183:2125–31.

30. Kotlar HK, Lewin A, Johansen J et al. High coverage sequencing of DNA from microorganisms living in an oil reservoir 2.5 kilometres subsurface. Environmental Microbiology Reports 2011;3:674–81.

31. Lazar I, Petrisor I, Yen T. Microbial enhanced oil recovery (MEOR). Petrol Sci Tech 2007;25:1353–66.

32. Lin J, Hao B, Cao G et al. A study on the microbial community structure in oil reservoirs developed by water flooding. J Petrol Sci Eng 2014;122:354–9.

33. Lozupone C, Knight R. UniFrac: a new phylogenetic method for comparing microbial communities. Appl Environ Microbiol 2005;71:8228–35.

34. McDonald D, Price MN, Goodrich J et al. An improved Greengenes taxonomy with explicit ranks for ecological and evolutionary analyses of bacteria and archaea. Isme J 2012;6:610–8.

35. Meyer M, Stenzel U, Hofreiter M. Parallel tagged sequencing on the 454 platform. Nature Protocols 2008;3:267–78.

36. Nishanthi R, Kumaran S, Palani P et al. Screening of biosurfactants from hydrocarbon degrading bacteria. J Ecobiotechnol 2010;2:47–53.

37. Olsson-Francis K, Pearson VK, Schofield PF et al. A study of the microbial community at the interface between granite bedrock and soil using a culture-independent and culture-dependent approach. Adv Microbiol 2016;6:233–45.

38. Orphan V, Taylor L, Hafenbradl D et al. Culture-dependent and culture-independent characterization of microbial assemblages associated with high-temperature petroleum reservoirs. Appl Environ Microbiol 2000;66:700–11.

39. Pantazaki AA, Dimopoulou MI, Simou OM et al. Sunflower seed oil and oleic acid utilization for the production of rhamnolipids by Thermus thermophilus HB8. Appl Microbiol Biotechnol 2010;88:939–51.

40. Parks DH, Tyson GW, Hugenholtz P et al. STAMP: statistical analysis of taxonomic and functional profiles. Bioinformatics 2014;30:3123–4.

41. Portwood J. A commercial microbial enhanced oil recovery technology: evaluation of 322 projects. SPE J 1995; DOI: 10.2118/29518-MS.

42. Ren HY, Zhang XJ, Song Z et al. Comparison of microbial community compositions of injection and production well samples in a long-term water-flooded petroleum reservoir. Plos one 2011;6, DOI: 10.1371/journal.pone.0023258.

43. Sen R. Biotechnology in petroleum recovery: The microbial EOR. Progr Energ Combust Sci 2008;34:714–24.

44. Shibulal B, Al-Bahry SN, Al-Wahaibi YM et al. Microbial enhanced heavy oil recovery by the aid of inhabitant spore-forming bacteria: an insight review. The Scientific World Journal 2014; DOI: 10.1155/2014/309159.

45. Wang LY, Duan RY, Liu JF et al. Molecular analysis of the microbial community structures in water-flooding petroleum reservoirs with different temperatures. Biogeosciences 2012;9:4645–59.

46. Wang LY, Ke WJ, Sun XB et al. Comparison of bacterial community in aqueous and oil phases of water-flooded petroleum reservoirs using pyrosequencing and clone library approaches. Appl Microbiol Biotechnol 2014;98:4209–21.

47. Youssef N, Elshahed MS, McInerney MJ. Chapter 6 Microbial processes in oil fields: culprits, problems, and opportunities. Adv Appl Microbiol 2009;66:141–251.

48. Youssef N, Simpson D, Duncan K et al. In situ biosurfactant production by Bacillus strains injected into a limestone petroleum reservoir. Appl Environ Microbiol 2007;73:1239–47.

49. Zhang F, She YH, Chai LJ et al. Microbial diversity in long-term water-flooded oil reservoirs with different in situ temperatures in China. Scientific Reports 2012;2:760.

50. Zhou J, Bruns MA, Tiedje JM. DNA recovery from soils of diverse composition. Appl Environ Microbiol 1996;62:316–22.

51. Zou C, Wang M, Xing Y et al. Characterization and optimization of biosurfactants produced by Acinetobacter baylyi ZJ2 isolated from crude oil-contaminated soil sample toward microbial enhanced oil recovery applications. Biochem Eng J 2014;90:49–58.

